# Genetic manipulation of candidate phyla radiation bacteria provides functional insights into microbial dark matter

**DOI:** 10.1101/2023.05.02.539146

**Authors:** Yaxi Wang, Larry A. Gallagher, Pia A. Andrade, Andi Liu, Ian R. Humphreys, Serdar Turkarslan, Kevin J. Cutler, Mario L. Arrieta-Ortiz, Yaqiao Li, Matthew C. Radey, Jeffrey S. McLean, Qian Cong, David Baker, Nitin S. Baliga, S. Brook Peterson, Joseph D. Mougous

## Abstract

The study of bacteria has yielded fundamental insights into cellular biology and physiology, biotechnological advances and many therapeutics. Yet due to a lack of suitable tools, the significant portion of bacterial diversity held within the candidate phyla radiation (CPR) remains inaccessible to such pursuits. Here we show that CPR bacteria belonging to the phylum Saccharibacteria exhibit natural competence. We exploit this property to develop methods for their genetic manipulation, including the insertion of heterologous sequences and the construction of targeted gene deletions. Imaging of fluorescent protein-labeled Saccharibacteria provides high spatiotemporal resolution of phenomena accompanying epibiotic growth and a transposon insertion sequencing genome-wide screen reveals the contribution of enigmatic Saccharibacterial genes to growth on their Actinobacteria hosts. Finally, we leverage metagenomic data to provide cutting-edge protein structure-based bioinformatic resources that support the strain *Southlakia epibionticum* and its corresponding host, *Actinomyces israelii*, as a model system for unlocking the molecular underpinnings of the epibiotic lifestyle.

## Introduction

The vast majority of metagenomic DNA sequences obtained from microbial species-rich environmental sources is derived from Bacteria and Archaea that have not been cultivated.

Conservative estimates suggest that these sequences, often referred to as microbial dark matter, represent organisms constituting approximately half of phylum level diversity within these domains^1^. Microbial dark matter holds great interest as a reservoir of biosynthetic pathways and enzymes with potential for biotechnological application^2^. In addition, understanding the functions of these genes is paramount to defining the molecular processes supporting a given ecosystem and for unraveling the physiology and cell biology of the organisms within^3^.

The candidate phyla radiation (CPR) of bacteria are reported to contain well over 50 phyla, only four of which have cultivated representatives^4^. This large monophyletic lineage of phyla and superphyla, including the group currently known as Patescibacteria, is thought to represent 15-50% of total bacterial diversity and contributes disproportionately to microbial dark matter^5^. Common features of CPR bacteria are their small size (as little as 100-200 nm in width), reduced genomes (typically < 1 megabase), and limited metabolic capability^6^. This has led to the hypothesis that these organisms broadly share a requirement for host organisms to support their growth. Indeed, experiments show that most cultivated CPR bacteria attach to and proliferate on the surface of other bacteria – living as obligate epibionts^4 7–10^. Genomic analyses have yielded speculation regarding the molecular functions that support the epibiotic lifestyles of CPR bacteria^5, 11^. However, owing to the phylogenetic distance separating CPR and well characterized bacteria, the function of much of their proteome cannot be predicted and a lack of genetic tools for these organisms has heretofore precluded the experimental investigation of genotype– phenotype relationships.

Among the CPR, members of the phylum Saccharibacteria, originally named Tm7, were the first to be cultivated in the laboratory^8^. Saccharibacteria are found in a multitude of terrestrial and marine environments, yet early interest in them stemmed from their widespread occurrence in human oral microbiomes^12^. Archaeological findings show this association dates to before the mesolithic period and recent work links Saccharibacteria to human oral health^12–14^. The growth of Saccharibacteria relies on the co-cultivation of host bacteria belonging to the class Actinomycetia with the phylum Actinomycetota, for which they exhibit strain level specificity^15–17^. Employing panels of Actinomycetia strains for Saccharibacterial enrichment has facilitated the isolation and sequencing dozens of strains^16, 18, 19^. Despite this progress, the phylum remains poorly sampled, with many divergent clades uncultivated, and the extent of genetic diversity unresolved.

The genome sequences of Saccharibacteria reveal common features that provide insight into molecular functions underlying their cellular physiology and lifestyle. Akin to other CPR bacteria, Saccharibacteria generally lack a respiratory chain, and pathways for the *de novo* generation of amino acids, nucleotides, and fatty acids^6^. On the contrary, Saccharibacteria universally possess a relative wealth of specialized secretory mechanisms including the type II and IV secretion systems (T2SS, T4SS)^11^. These diverge significantly from related systems in bacterial pathogens that deliver toxins and effectors to eukaryotic host cells; however, it has been speculated that they function in an analogous fashion to support bacterial host co-option by Saccharibacteria^11^. Saccharibacteria also possess type IV pili (T4P), which were implicated in twitching motility and host adhesion through the use of a small molecule inhibitor of pilus extrusion^17^. Though such genomic analyses and experiments provide fertile ground for the formulation of hypotheses, progress toward a mechanistic understanding of the unique biology of Saccharibacteria and CPR bacteria as a whole has been stymied by a lack of genetic tools^20^. Here, we discover that natural competence can be harnessed for genetic manipulation of Saccharibacteria. With this capability in hand, we go on to use fluorescent protein expression to conduct time-lapse microscopic analysis of the Saccharibacterial lifecycle and we perform transposon mutagenesis to identify genes important for epibiotic growth. Our findings offer an initial mechanistic glimpse into the cellular functions encoded in microbial dark matter.

## Results

### Isolation and characterization of Saccharibacteria strains

We collected, pooled, homogenized, and filtered saliva and dental plaque from volunteers, and enriched this material for Saccharibacteria using the method described by Bor *et al*^18^. This led to the isolation of two strains, which exhibit distinct host specificity (Figure 1A). Phylogenetic analysis using concatenated alignments of 49 core, conserved protein sequences obtained through complete genome sequencing placed the two strains within human oral subclades of the G1 clade of Saccharibacteria (Figure 1B-E)^21^. Based on these assignments, we named our strains *Candidatus Nanosynbacter lyticus* ML1 (*Nl*) and *Ca. Southlakia epibionticum* ML1 (*Se*). The genome sequences of the two strains additionally indicated they bear features typical of Saccharibacteria, such as genes associated with specialized secretion systems, cell surface appendages, and competence, coupled with a lack of genes required for a multitude of biosynthetic pathways and an overall reduced genome size (Figure 1D,E).

**Figure 1.**
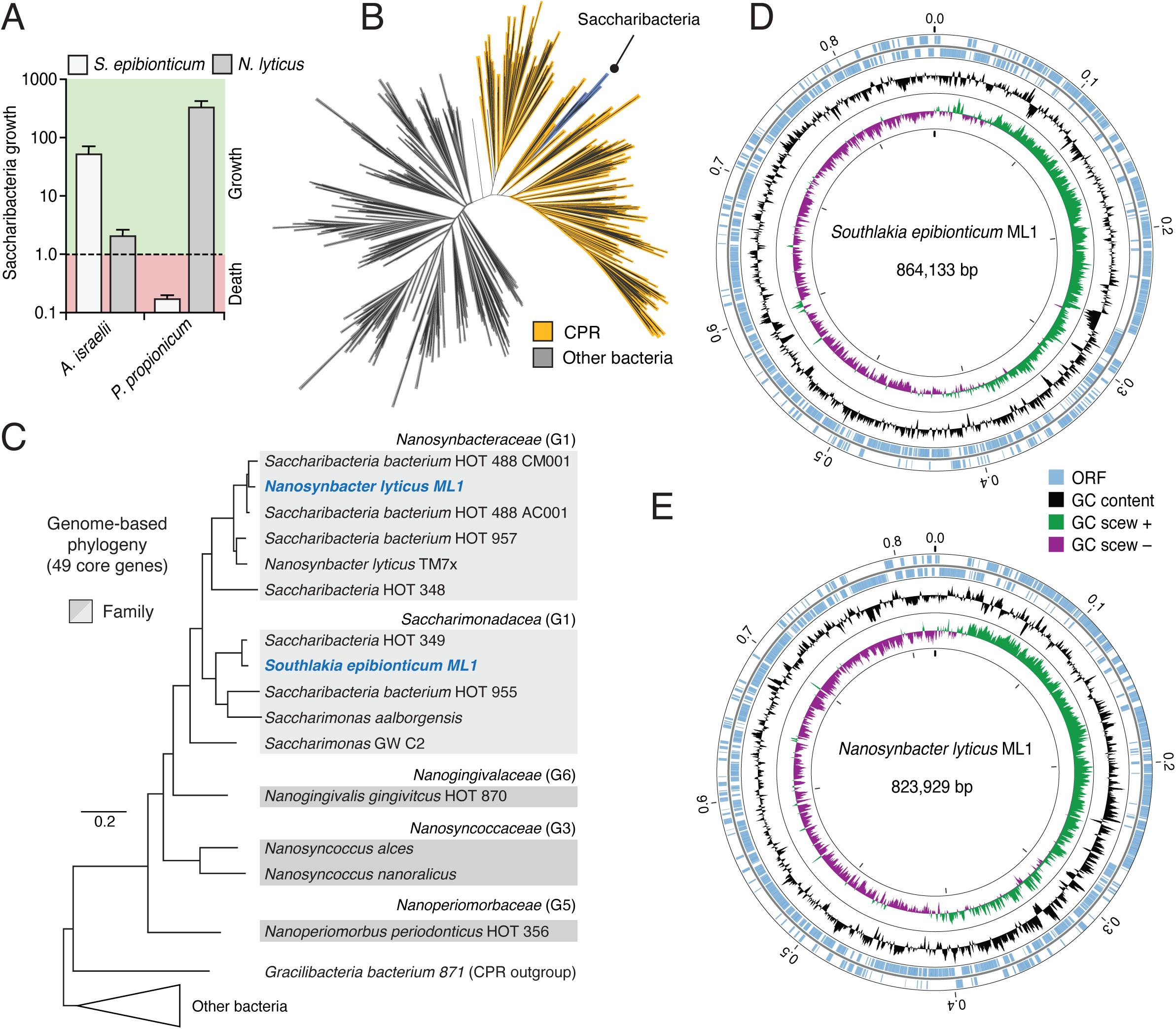
Phylogenetic placement and genome sequencing of newly isolated Saccharibacteria strains *S. epibionticum* ML1 (*Se*) and *N. lyticus* ML1 (*Nl*). (A) Maximum growth (fold change) achieved by *Se* and *N*l during co-culture with compatible host species *A. israelii* and *Propionibacterium propionicum* respectively, and population change (growth or death) detected at equivalent timepoints with an incompatible host. (B) Phylogeny of CPR and other bacteria based on concatenated ribosomal proteins, with the placement of the Saccharibacteria phylum indicated. Figure adapted from Castelle *et al.*^6^ (C) Phylogeny constructed using 49 core, universal genes indicating placement of *Se* and *N. l* (blue text) within Saccharibacteria. Family names (as designated by the Genome Taxonomy Database) and groups previously designated by McLean *et al.* (G1, etc.) are indicated for each clade^11^. HOT, human oral taxon. (D, E) Overview of the genome sequences of *S. epibionticum* ML1 and *N. lyticus* ML1.

### Genetic manipulation of Saccharibacteria via natural transformation

We sought to develop methods for reverse genetic analyses within Saccharibacteria. Genes encoding the core components of the *com* DNA uptake system are conserved across CPR phyla, including the Saccharibacteria^5^. These consist of ComEC, the central membrane DNA conduit, DprA, a catalyst of Rec-mediated recombination, and ComFC, which binds ssDNA and links import to recombination^22–24^. Com proteins function in concert with type IV pili, which are also widely distributed in Saccharibacteria, and CPR bacteria more generally^5, 23^. Given the lack of nucleotide biosynthetic capability in CPR bacteria, it has been proposed that these systems facilitate nucleotide acquisition^5^. Although the presence of the Com system is largely not predictive of DNA transformation in a laboratory setting^25^, we sought to test whether exogenous DNA could be exploited for genetically manipulating Saccharibacteria.

As a first step toward assessing feasibility of genetics in Saccharibacteria, we searched for antibiotics with convenient resistance determinants that potently inhibit the growth of *Se* without impacting that of its preferred host, *Actinomyces israelii* F0345 (*Ai*). These experiments revealed that across a wide range of concentrations, the aminoglycoside hygromycin fulfills these criteria (Figure S1). Next, we designed and generated a linear cassette containing the hygromycin resistance gene (*hph*) codon optimized for *Se* flanked by the promoter and terminator regions of the TM7x *tuf* gene (elongation factor Tu), an open reading frame (ORF) predicted to be highly expressed (Figure 2A). For the insertion of this cassette, we selected an intergenic region located between two convergently transcribed ORFs, SEML1_0215 and SEML1_0216, hereafter referred to as neutral site 1 (NS1). To promote homologous recombination, approximately 1000 bp on either side of the insertion site were added to the 5′ and 3′ ends of our cassette.

**Figure 2.**
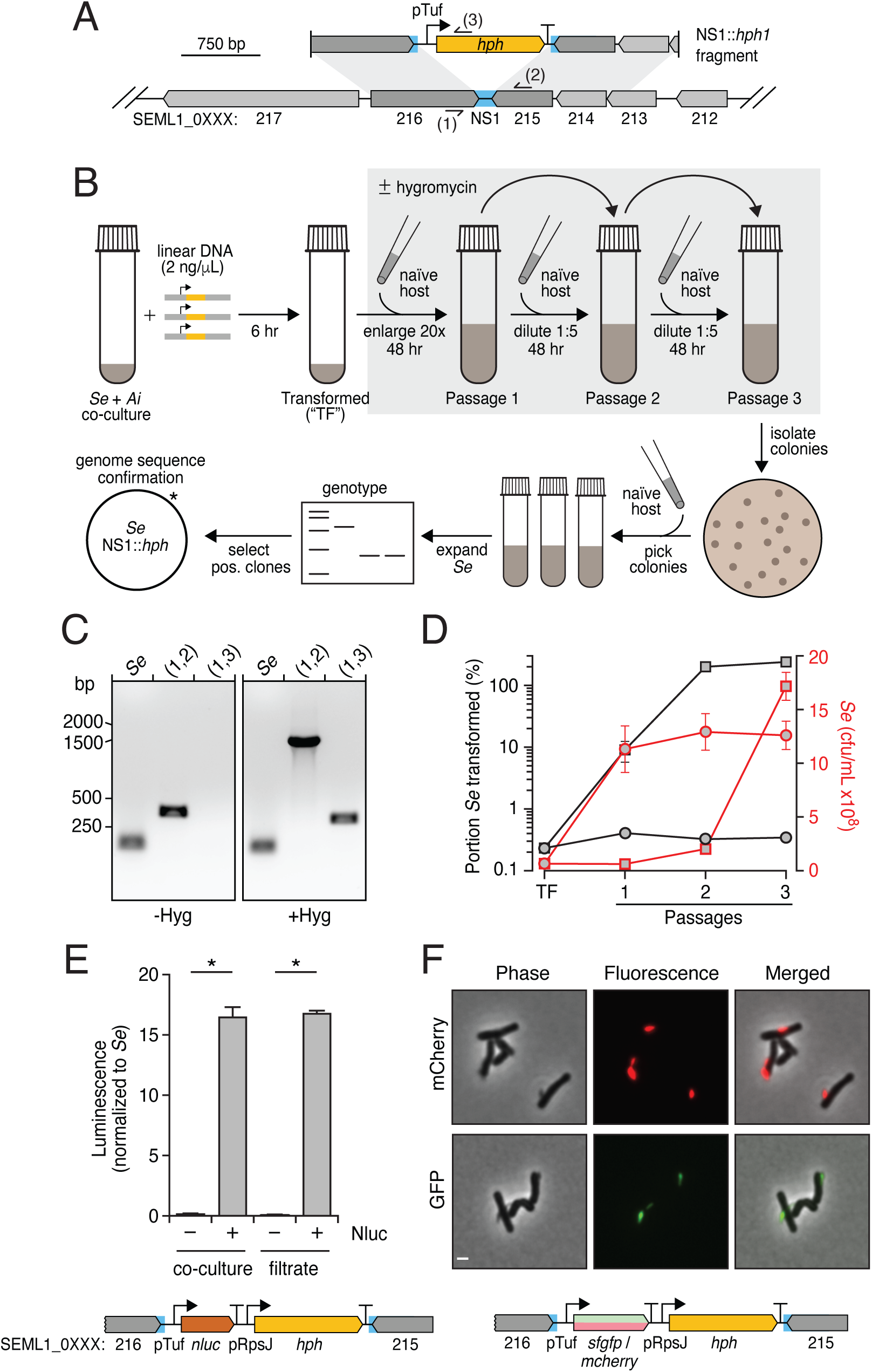
Harnessing natural transformation to generate mutant Saccharibacteria. (A) Schematic depicting the intergenic neutral site (blue, NS1) targeted for insertion of a hygomycin resistance cassette (yellow) in the *Se* genome and the linear DNA fragment employed in transformation experiments. Primer binding sites used for genotyping are indicated (sites 1-3). (B) Overview of the *Se* transformation protocol. After incubation with linear DNA, *Se + Ai* co-cultures are enlarged concomitant with hygromycin addition and serially passage with addition of naïve host at each dilution to promote *Se* growth (grey box). Clonal transformed *Se* populations were obtained by plating to isolate single colonies of *Ai* with accompanying *Se* cells, followed by growth in liquid culture, with additional *Ai,* to promote *Se* population expansion. (C) PCR-based genotyping of *Se* clones obtained following transformation according to the protocol show in (B) in the presence (right) or absence (left) of selection with hygromycin during the expansion and passaging steps. Binding sites for primers targeting NS1 (1, 2) and *hph* (3) are shown in (A). Positive control primers (*Se*) target a locus distant from NS1. (D) Growth (red) and percent of *Se* transformed (grey) over the course of transformation protocol depicted in (B), in the presence (squares) or absence (circles) of selection with hygromycin. (E) Luminescence production from *Se–Ai* co-cultures (left) or co-culture filtrates (right) in which *Se* contains a nanoluciferase expression cassette inserted at NS1 (shown at bottom). (F) Fluorescence and phase contrast micrographs of *Se–Ai* co-cultures in which *Se* carries an *mcherry* (top) or *sfgfp* (bottom) expression cassette inserted at NS1. See also Figure S2. Data in (E) represent mean ± s.d. Asterisks indicate statistically significant differences (unpaired two-tailed student’s t-test; *p<0.05).

To transform *Se*, we incubated *Se*–*Ai* co-cultures with 2.0 ng/μL of our linear cassette for six hours before initiating selection with hygromycin (Figure 2B). Naïve host was added concomitantly in order to permit the outgrowth of successfully transformed *Se* cells. Reasoning that transformation may be inefficient, we passaged these co-cultures twice, at 48 hour intervals, with continued hygromycin selection and the addition of naïve host. Cultures were then diluted and plated to obtain colonies, where were selected and propagated with naïve host without selection before genotyping (Figure 2B). The latter step was included in order to bottleneck the *Se* population and facilitate the isolation of clonal populations. Remarkably, each *Se*-infected culture we tested – accounting for the majority of colonies selected – contained our synthetic cassette inserted at the expected location (Figure 2C). Whole genome sequencing confirmed these PCR results and it further showed that cassette integration occurred without introducing off-target mutations.

Quantitative PCR (qPCR) analysis of total *Se*, transformed *Se*, and *Ai* at regular intervals during our transformation procedure demonstrated that approximately 0.2% of Se contain the integrated cassette by the conclusion of the initial incubation period (Figure 2D). Though *Se* levels remain low through the second passage under selection with hygromycin, all surviving *Se* cells bear the cassette at this time point. In the final passage, the population of *Se* continues to maintain the cassette and expands markedly, far surpassing levels of the host (Figure S2A). In the absence of hygromycin, similar quantities of initially transformed *Se* are observed; however, this small proportion fails to expand despite overall robust growth of *Se*. We observed similar transformation behavior using lengths of DNA with homology to the insertion site flanking regions as short as 221 bp (Figure S2A) and with as little as 0.02 ng/μL (Figure S2B). Finally, to probe the generality of our methods, we identified a predicted neutral site within *Nl* and subjected this second Saccharibacterial strain to an analogous transformation protocol. Genotyping of transformed populations indicated cassette insertion at the desired location also occurred within this strain (Figure S2C,D).

The ability to introduce heterologous DNA into *Se* has numerous foreseeable applications, one of which is the expression of reporter genes that allow *Se* to be distinguished and studied within the context of co-culture with their hosts. To explore this possibility, we designed and generated NS1 insertion cassettes containing genes encoding nanoluciferase, mCherry and GFP under the control of the *tuf* promoter and upstream of *hph* driven by a second predicted strong promoter of TM7x, that of *rpsJ* (Figure 2E,F). Using our transformation protocol, we obtained clonal populations of hygromycin-resistant *Se* containing each of these genes. Luminescence assays and fluorescence microscopy demonstrated robust activity of each reporter gene (Figure 2E,F). We did not detect their activity in host cells, indicating the feasibility of achieving specific manipulation of *Se* in the context of a co-culture.

The expression of fluorescent proteins within *Se* provided the opportunity to visualize CPR bacterial growth with extended, time-lapse fluorescence imaging. Over the course of 20 hrs, both *Se* and *Ai* populations – deposited from co-cultures at low *Se*:*Ai* ratio onto an agar substrate containing growth media – showed clear evidence of expansion (Videos [S1-11](https://youtube.com/playlist?list=PLSQI8BpslH-PhSKoxXLm7Ur8zJneoS5We)). Apparent T4P-mediated motility of *Se* was also observed, as reported by Xie *et al*^17^. Our time-lapse imaging further captured features of the CPR lifecycle at unprecedented spatiotemporal resolution. For instance, we could distinguish productive (*Se* growth supporting) versus non-productive (*Se* adhered without concomitant growth) interactions, and directly measure their respective impact on individual host cells (Figure 3). Additionally, we observed productively adhered mother cells producing small swarmer cell progeny via repeated polar budding, and the differentiation of a subset of these progeny into mother cells (Videos [S1-11](https://youtube.com/playlist?list=PLSQI8BpslH-PhSKoxXLm7Ur8zJneoS5We)). Altogether, these findings show that natural transformation can be exploited to genetically manipulate CPR bacteria in a directed manner and open a window into the distinctive biology of this largely unexplored group of organisms.

**Figure 3.**
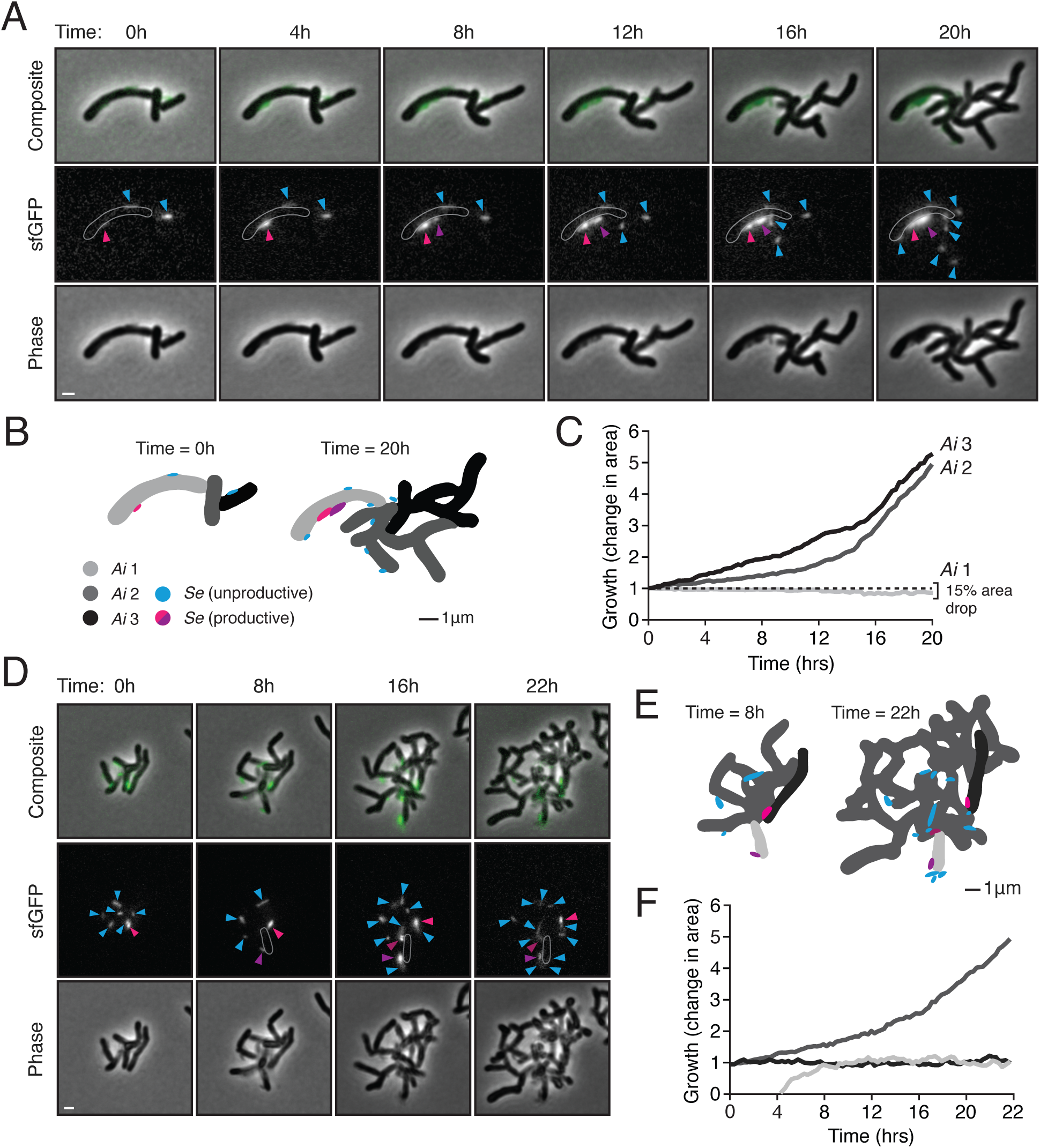
Fluorescent protein expression and quantitative microscopy enable tracking of the *S. epibionticum* lifecycle. (A) and (D) Snapshots captured at the indicated time points from timelapse fluorescence and phase contrast microscopy of GFP-expressing *Se* grown in co-culture with *Ai.* Arrows indicate *Se* cells exhibiting productive (pink, purple) and non-productive (blue) interactions with *Ai* cells. White outlines in the fluorescent channel indicate depict an *Ai* cell affected by *Se* infection (*Ai* 1). (B) and (E) Omnipose-generated segmentation of *Se* and *Ai* cells depicted in (A) and (D), at the start (left) and end (right) of the 20 or 22 hr growth period. (C) and (F) Growth of individual *Ai* cells as impacted by productive (light grey) or non-productive *Se* cells (black, dark grey). Colors correspond to cell masks shown in (B) and (E). For the full time course captured in (A) and (D), see Videos [S1](https://www.youtube.com/playlist?list=PLSQI8BpslH-PhSKoxXLm7Ur8zJneoS5We) and [S2](https://www.youtube.com/playlist?list=PLSQI8BpslH-PhSKoxXLm7Ur8zJneoS5We), respectively. For additional examples of tracked *Se–Ai* growth, see Videos [S3-S11](https://youtube.com/playlist?list=PLSQI8BpslH-PhSKoxXLm7Ur8zJneoS5We).

### Tn-seq of *Se* provides insights into the epibiotic lifestyle

The ability to genetically manipulate Saccharibacteria enables myriad avenues of investigation. As a first step toward genetic dissection of the *Se* epibiotic relationship with *Ai*, we conducted transposon-insertion sequencing (Tn-seq) within *Se* during growth on *Ai*. To this end, we performed *in vitro* Tn5-based transposition on purified *Se* genomic DNA, repaired gaps as described by Manoil and colleagues, and used sequencing to confirm high frequency, homogenous insertion across the genome. This DNA was then used to transform *Se*, with slight modifications from our basic transformation protocol (see Methods). Most notably, we elected to increase the scale of the experiment to account for our previously measured transformation efficiency and thereby avoid population bottlenecking following the onset of selection. We collected four samples for Tn-seq analysis, an initial sample following the transformation and recovery period (T0), and three additional samples representing the population after 48 hr serial passages with the addition of naïve host (T1-3). Measurements of *Se* and *Ai* levels at each of these timepoints revealed a precipitous drop in *Se* levels at T1 that is not observed at a similar timepoint in transformations targeting NS1, suggesting that the large majority of transposon insertions in *Se* are deleterious (Figure S3). Our measurements also showed that, as expected, *Ai* levels drop relative to those of *Se* at later passages, such that by T3, *Se* outnumbers *Ai* by approximately 50-fold.

Preliminary analysis of our Tn-seq data indicated that the T0 population of *Se* contained a large number of unique insertions (9,996) relative to subsequent timepoints (T1, 1709; T2, 1887; T3, 1699) despite similar sequencing depth (Table S1, Figure 4A). Furthermore, application of the TRANSIT algorithm to the T0 data identified a small number of essential genes relative to our expectations (905 *Se* genes; 56 essential by Gumbel or HMM)^26^. Taken together with our measurement of *Se* and *Ai* population levels, we interpret these findings as evidence that little selection occurred prior to T0 sampling. Therefore, we compared subsequent timepoints to T0 using TRANSIT resampling in order to assess gene level fitness contributions throughout the experiment. In total, this approach identified 214 genes that demonstrate statistically significant (multiple-comparison adjusted P value < 0.05) fitness contributions across all timepoints, with selection yielding greater numbers at each subsequent sampling (T1, 222; T2, 252; T3, 275) (Table S1).

**Figure 4.**
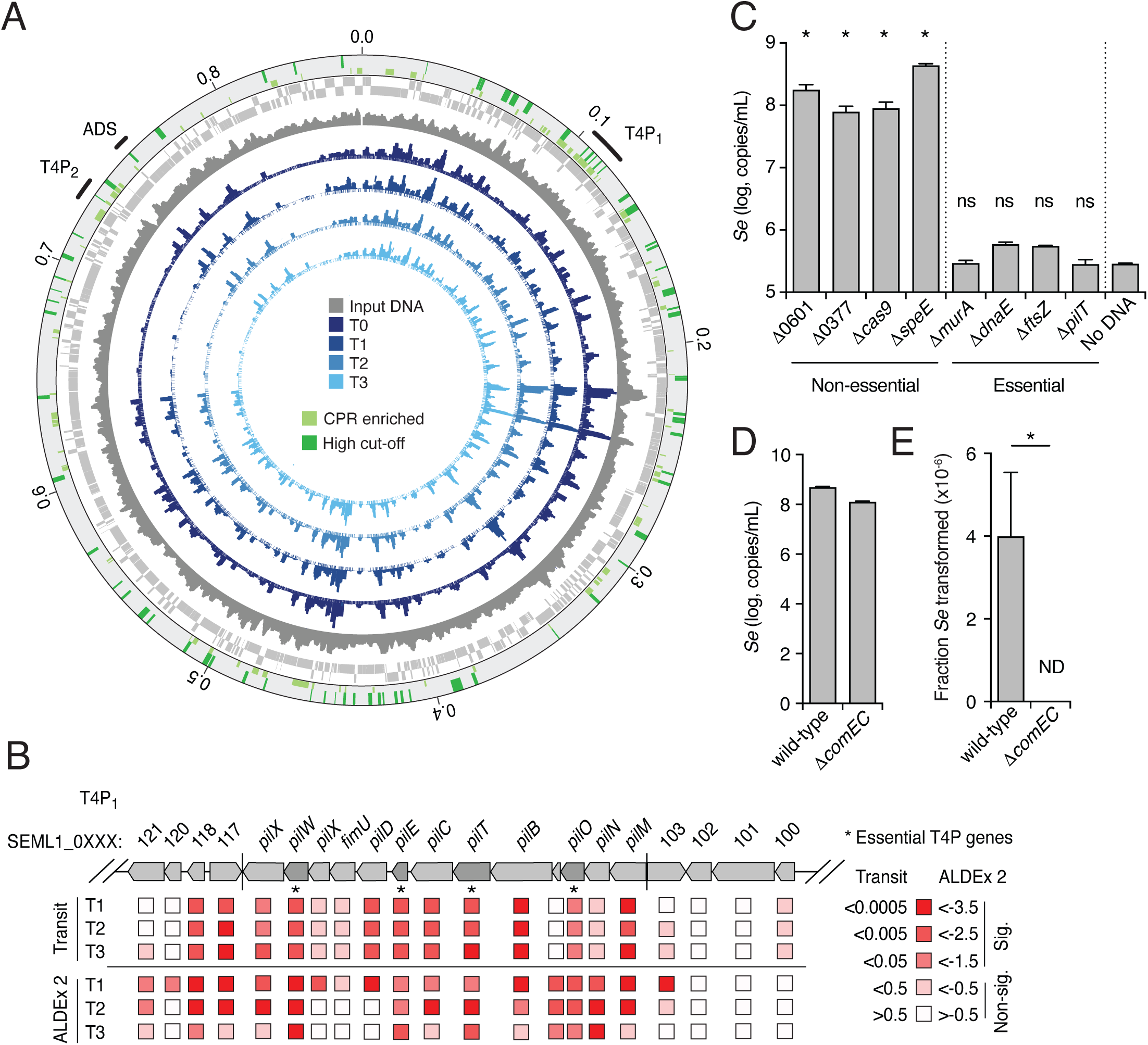
Identification of genes important for fitness of *S. epibionticum* during co-culture with *Ai* identified by Tn-seq. (A) Overview of normalized transposon insertion frequency across the *Se* genome detected in input DNA used for mutagenesis (dark grey), and from samples collected immediately following transformation (T0, dark blue) and subsequent outgrowth time points (T1-T3, shades of blue). Genes encoding proteins belonging to CPR-enriched protein families (light green) and those found to be significantly important for fitness across all time points and by two different metrics (dark green). The location of the arginine deiminase system (ADS) genes and two loci containing T4P genes (T4P1, T4P2) are indicated in the outer circle. (B) Schematic depicting T4P1 and flanking genes (separated by vertical lines) and the relative fitness contribution of each gene as determined by the TRANSIT resampling algorithm (adjusted p- values) and ALDEx2 (*delta* scores). Asterisks indicated genes found to be significant by both metrics across all time points. Gene annotations were derived from Foldseek queries using AF models generated for each gene product (see methods). (C) *Se* population levels detected in *Se– Ai* cocultures following transformation with constructs designed to replace the indicated genes with *hph.* Candidate essential genes tested were considered significant for *Se* fitness by TRANSIT and ALDEx2 at a minimum of 2 of 3 time points (see Table S1). (D, E) Total *Se* population (D) and proportion transformed (E) following transformation with an unmarked cassette targeted to NS1 in the indicated strains of *Se.* See also Figures S3 and S4, Table S1. Data in (C-E) represent mean ± s.d. Asterisks indicate statistically significant differences (C, one-way ANOVA followed by Dunnett’s compared to no DNA control; E, unpaired two-tailed student’s t-test; *p<0.05, ns, not significant).

The transposon insertion density we observed in our T1-3 samples is below the value needed to obtain optimal results in TRANSIT. Given this potential caveat, we elected to further process our data with ALDEx2, a program widely used for differential abundance analysis of compositional data^27^. Overlaying genes with significantly differential insertion abundance determined by ALDEx2 with those we obtained using TRANSIT revealed a group of 79 genes that we deem unambiguously critical for *Se* growth on *Ai* in the context of our experimental conditions (Table S1 and Figure 4A). This group includes genes encoding proteins that comprise core cellular machinery such as DNA polymerase, RNA polymerase and the ribosome.

Interestingly, the group also includes non-canonical essential genes that likely encode specialized functions that support the epibiotic lifestyle of CPR bacteria; examples include genes within a T4P operon and several genes lacking homologs in non-CPR bacteria (Figure 4B).

If our Tn-seq findings provide an accurate measure of gene-level fitness contributions, then those identified as critical for *Se* fitness via TRANSIT or TRANSIT and ALDEx2 should be difficult to inactivate by targeted mutagenesis. Accordingly, to validate these data, we selected four genes identified as significant contributors to fitness in one or both analyses, along with four control genes, and designed constructs to replace each with an *hph* expression cassette. The constructs shared equivalent length flanking sequences to enable the direct comparison of their behavior in our transformation protocol. We assessed the fitness impact of inactivating each gene by quantifying *Se* populations following transformation and outgrowth under selection with hygromycin. For the control genes not predicted to contribute to fitness, we achieved robust levels of *Se* growth by this time point, comparable to that achieved when introducing the *hph* expression cassette at NS1 (Figure 4C). In contrast, in transformations targeting each of the four predicted essential genes examined, *Se* failed to proliferate, consistent with the inactivation of these genes strongly impacting fitness. Together, these findings provide confirmation that our Tn-seq analysis successfully identified the relative fitness contributions of *Se* genes during co-culture with *Ai*.

We noted that genes encoding homologs of *com* system components ComEC, ComF and DprA were not among those genes defined as contributing significantly to *Se* fitness in our Tn-seq study. This finding suggests that this DNA competence machinery is not required for acquiring nucleotides to support *Se* growth, as had been previously suggested^5^. To determine whether the *com* system of *Se* instead functions to mediate natural transformation, we generated an *Se* strain in which the *comEC* ORF is replaced by the *hph* expression cassette (*Se* /-*comEC*::*hph*). To assess whether this mutation affects *Se* transformation, we measured the efficiency of inserting a second, unmarked cassette at NS1. At the conclusion of the transformation protocol, insertion at the NS1 site was only detectable in the wild-type background, supporting the hypothesis that the *com* system mediates natural transformation in this species (Figure 4D,E).

Banfield and colleagues previously leveraged the large number of CPR genomes available from metagenomic and traditional genome sequencing datasets (n=2,321) to define CPR-enriched protein families^5^. To gain insight into which of these contribute to fitness during *Se* growth on *Ai*, we identified *Se* proteins belonging to these families, and cross-referenced hits against the gene lists obtained from our Tn-seq experiment following TRANSIT or TRANSIT and ALDEx2 analysis (Figure S4). Interestingly, we found that the representation of CPR- enriched protein families among proteins critical for *Se* survival on *Ai*, 22% and 19%, respectively, is considerably higher than their proportion within the overall *Se* proteome (12%) (Table S1 and Figure S4). As many of these protein families have no predicted function, nor orthologs outside of CPR bacteria, their future study is likely to reveal unique features of these organisms.

### Development of resources for dissecting Saccharibacteria–host cell interactions

With available genetic tools and gene-level fitness data, *Se* could become a tractable model for the study of Saccharibacteria. However, comprehensive genotype–phenotype dissection of the interaction between *Se* and its host remains limited by lack of equivalent tools and data for *Ai.* As a first step towards bridging this gap, we obtained a complete genome sequence for the host strain we employed, *A. israelii* F0345 (Figure S5A). This strain was isolated from the human oral cavity as part of the Human Microbiome Project, but a genome sequence had not been previously deposited in a public database. Although genetic tools have not been developed for this organism, gene inactivation via the electroporation-mediated introduction of suicide plasmid or linear gene inactivation constructs has been achieved in related species^28, 29^. The availability of genetic information for this strain should facilitate development of selectable markers and counter selection strategies.

Automated annotation of the *Se* genome sequence failed to predict functions for the proteins encoded by many ORFs (278 of 855), including 14/79 belonging to the conservative list of essential genes we identified by Tn-seq (Table S1). To improve functional predictions associated with ORFs in the *Se* genome, we applied a battery of sequence and structure-based computational tools. ProtNLM is a natural language model trained on UniProt to predict protein names given a sequence^30^. Applying ProtNLM to the *Se* proteome yielded functional annotations for 337 of 855 proteins (ProtNLM score ≥ 0.5, Table S2). This sequence-based annotation was further improved by mapping each protein to Pfam domains using hmmscan, detecting Pfam domains in an additional 270 proteins (see Methods for cutoffs), 245 of which could be assigned functions^31, 32^ (Table S2).

To complement our sequence-based annotations, we took advantage of recent advances in protein structure prediction to conduct genome-wide structure-based homology analyses of the *Se* proteome^33, 34^. Generation of structural models using AlphaFold (AF) relies on evolutionary information extracted from multiple sequence alignments (MSAs)^34^. For nearly 25% of the *Se* proteome (220 proteins), our initial MSAs based on HHblits searches of UniRef and BFD (the default databases used by AF) were too shallow for high confidence structure prediction (<500 sequences post filtering, Figure 5A)^35–37^. To improve the MSA depth for these proteins, we implemented a hidden Markov model (HMM)-based approach to identify and align additional homologs for each protein from multiple metagenomic datasets. This resulted in considerably deeper MSAs (>500 sequences) for an additional 9% of the proteome (Figure 5B). Using the highest depth MSA obtained for each protein, we obtained AF models for >99% of the predicted *Se* proteome. For comparison, we also computed AF models using the original 220 MSAs that contained <500 sequences and compared the model confidence metric obtained using these and the improved depth MSAs. Particularly for low confidence models obtained using MSAs generated with the default databases (average pLDDT<50), we found that the use of the deeper MSAs for structural prediction resulted in substantial model confidence improvement (Figure 5C). The structure model improvements enabled by using extensive metagenomic databases for MSA generation were further underscored by structural homology search results obtained using Foldseek (FS)^38^. For some proteins, the improved structural models led to the identification of structural homologs for proteins that initially had no FS matches passing our cutoffs, while for others, the changes in the overall predicted structure led to the identification of different and more closely aligning top matches (Figure 5D,E, Figure S5B-D). In total, using FS we were able to identify similar structures for 88% of modeled proteins (748/852, Table S2).

**Figure 5.**
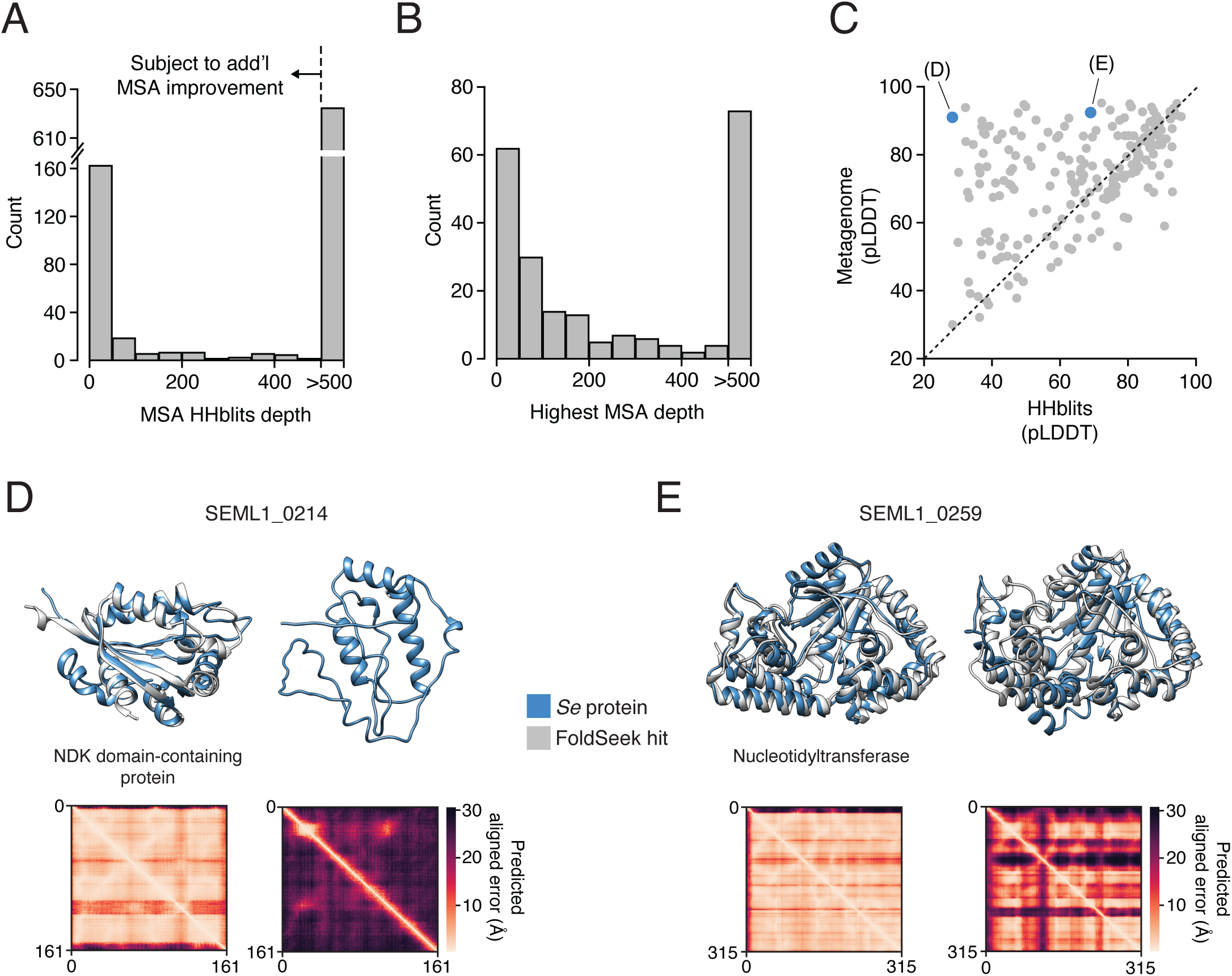
Inclusion of extensive metagenomic data in MSAs enables proteome-wide AF modeling of *Se* protein structures. (A) Histograms depicting MSA depths obtained for *Se* proteins using HHblits. (B) Maximum depths obtained for *Se* protein MSAs that initially contained <500 sequences. Additional sequences were sourced from metagenomic sequence databases and incorporated into MSAs using Jackhmmer or Phmmer (see Methods). (C) Comparison of the AF confidence metric (pLDDT) determined using Hhblits or Jackhmmer/Phmmer (Metagenome)-generated MSAs for *Se* proteins with initially shallow MSAs (<500). *Se* proteins shown in (D) and (E) are highlighted in blue. (D and E) Example *Se* protein structure models and associated predicted alignment matrices obtained using shallow (right) or metagenomic sequence-improved (left) MSAs. *Se* proteins models (blue) are aligned to models from top FoldSeek (FS) hits (light grey, A0A2H0BDY7 (D), A0A8B1YPH4, metagenome and A0A3D0YBM7, shallow, (E)), when available. The annotation in (D) derives from the best FS hit; in (E), the best FS hit is an uncharacterized protein. The function for this protein was assigned using DPAM. Some structures are trimmed to highlight the alignment. See also Figure S5, Table S2.

The incorporation of structural information resulted in functional predictions for 70 *Se* proteins that had none assigned by sequence-based approaches (Table S2). One example relevant to our interest in genetics within *Se* is the protein encoded by SEML1_0370, for which sequence-based annotations assigned no function. A FS search performed using the AF-generated model for this protein revealed it shares overall structural homology to and active site residues with the class II restriction endonuclease *PvuII*^39^. We confirmed using the Restriction Enzyme Database that SEML1_0370 does not share detectable primary sequence homology with characterized restriction endonucleases. Characterization of the sequence recognized by this and other *Se* restriction enzymes could lead to improvements in transformation efficiency, as DNA cassettes could be designed to exclude their target sequence and thus subvert restriction^40^.

Many of the AF models with similar structure identified for *Se* proteins are uncharacterized (12%), providing little information regarding function. Moreover, unrelated proteins may share similar structure due to convergence, and additional evidence is usually needed to assess the homologous relationships. Domain Parser for AF Models (DPAM) is a recently developed tool that parses structural domains from AF modeled structures, and integrates both structure and sequence searches to map protein domains to Evolutionary Classification of protein Domains (ECOD)^41, 42^. Using DPAM, we detected homologous ECOD domains for 80% of predicted *Se* proteins (Table S2). This included 39 proteins for which no function was assigned by other methods. In supplementary material accompanying this report, we provide alignments for each mapped *Se* protein domain, along with links to corresponding Protein Data Bank identifiers (PDB IDs) and ECOD classifications (https://conglab.swmed.edu/ECOD_Se/Se_ECOD.html). In total, these predictions provide a foundation for future use of the *Se*–*Ai* system as a model for understanding CPR bacteria.

## Discussion

Scientists have been aware of CPR bacteria in environmental samples for many years, yet our understanding of this group of organisms has lagged^4^. One major challenge is their apparent strict requirement for host bacteria, the identity of which cannot currently be determined a priori, thus adding significant complexity to CPR isolation^7, 18, 43, 44^. It is foreseeable that methods bringing to bear both experimental and computational approaches could provide predictions of host–epibiont associations in the future. Here we addressed a second impediment for studying CPR bacteria – a lack of genetic tools for their manipulation. Indeed, despite streamlined methods for isolating Saccharibacteria on host Actinobacteria that have been employed by several laboratories, this latter challenge has to-date prohibited a molecular dissection of their epibiotic lifestyle^16, 17, 44^. Our discovery that natural transformation can facilitate targeted mutagenesis in Saccharibacteria opens the door to direct interrogation of genotype–phenotype relationships within these CPR bacteria; however, challenges remain. The most fundamental hurdle is that factors required for the Saccharibacteria–host cell interaction are also required for viability. In analogous situations, researchers overcome this using assorted conditional inactivation strategies (e.g. modulated expression, temperature sensitivity, inducible degradation). While it is conceivable that such methods could be implemented in Saccharibacteria, a powerful approach in this system may involve exploiting host genetics to identify gene interactions that suppress otherwise lethal mutations in the epibiont.

The small genome of Saccharibacteria stands in contrast to the apparent complexity of their lifecycle. Xie *et al.* proposed a model for the epibiotic growth of Saccharibacteria that consists of four stages: T4P-mediated infection, growth, bud formation and asymmetric division^17^. Our time-lapse microscopy generally supports this model; however, it further revealed facets of the Saccharibacteria–host interaction not previously captured. While the aforementioned model implies uniform progression, our data suggest the lifecycle is more complex. We find that only a small subset of *Se* achieve productive infections, and that after an extended enlargement period, these cells rapidly bud a large number of highly motile swarmer cell progeny (>10 in a 20 hr period observed in many instances). Quantitative analyses of our data additionally permitted us to link productive and non-productive *Se* infections with corresponding host cell outcomes. We observed a striking inverse relationship between *Se* growth and that of *Ai*; cells productively infected with as few as one *Se* not only failed to divide, but diminished in size, whereas those infected unproductively by multiple *Se* readily proliferated.

Our microscopy observations suggests a potential division of labor within *Se* populations, wherein one subpopulation is devoted to reproduction, and a second, motile subpopulation searches for new compatible host cells. Operating under the assumption that the Saccharibacteria–Actinobacteria relationship is chiefly one of parasitism, which we maintain is not fully resolved in the literature, then the relevant prevailing evolutionary theory predicts that *Se* fitness is dependent on both the rate at which it causes new infections and the duration of those infections^45^. The reproductive strategy of *Se* appears consistent with this framework; release of progeny cells destined to infect other host cells tempers host cell burden and prolongs the infection, while concurrently maximizing *Se* reproduction. A similar outcome could be achieved if *Se* employed active mechanisms to block superinfection, reminiscent of those utilized by phage; however, swarmer cell production has the added advantage of increasing *Se* dispersal in the face of low host cell densities^46^. Interestingly, most nascent swarmer cells we observed do not themselves establish a productive infection within our 20 hr observation period, despite close proximity and apparent adherence to host cells. It may be that the time required to establish a productive interaction varies, and many of these cells would go on to become productive. It is also conceivable that infected host cells defend against secondary infections, perhaps even inducing a defensive state in neighboring kin cells. Future studies coupling quantitative microscopy with genetics to further dissect the *Se*–*Ai* interaction will no doubt address these questions and shed light on this fascinating interphyla relationship.

Pili are a basal feature of CPR bacteria and our Tn-seq results highlight the critical role they play for these organisms^7, 17, 47^. Among the 15 genes that are both enriched in CPR and that we conservatively deemed essential in *Se*, six are predicted T4P components (Table S1). A prior study utilizing quercetin, a small molecule inhibitor of pilus retraction, reported that T4P are important for host attachment and twitching motility in a second Saccharibacterial species^17^. In naturally competent bacteria, T4P work in concert with the *com* system to bind and mediate the uptake of extracellular DNA^48, 49^. Our observation that *Se* cells lacking *comEC* are viable suggests that the essentiality of T4P is unrelated to its role in competence. *Se* lacks *de novo* nucleotide synthesis pathways, thus the organism presumably possesses an essential DNA or nucleotide uptake pathway that operates independently of the *com* system. Whether T4P participate in such an alternative DNA uptake mechanism remains unknown.

The arginine deiminase system (ADS) is an ATP generating catabolic pathway prevalent in mammalian-adapted Saccharibacteria species^50^. A prior study demonstrated that arginine supplementation to *N. lyticus* TM7x–*A. odontolyticus* co-cultures permits acid neutralization via ammonia production, supporting the viability of both strains during acid stress^50^. The authors of this study put forth a model in which hosts lacking ADS benefit from and are specifically permissive to colonization by Saccharibacteria containing the pathway. In *Se*, arginine deiminase (ArcA) is a fusion of two proteins, ArcA and ornithine carbamoyltransferase (ArcB); *arcA* is among our conservative list of genes deemed essential for *Se* growth. Inactivation of genes encoding the remaining ADS components, carbamate kinase ArcC and arginine/ornithine antiporter ArcDE, also yielded significant *Se* fitness defects by one or more metric (Table S1). Interestingly, in contrast to the proposed model for ADS function in Saccharibacterial–host cell interactions, we find intact homologs of each ADS gene in *Ai*. Taken together with our Tn-seq data, our results suggest that in some co-culture pairs, the organisms may compete for arginine. However, the mechanism by which the pathway influences Saccharibacteria–host cell dynamics *in vivo*, including specific niches in the oral cavity, remains an open question^50^.

We anticipate that the methods and genetic tools presented here will facilitate molecular level characterization of Saccharibacteria and CPR bacteria more broadly. In this regard, one key question is the extent to which natural competence is active and able to be similarly exploited across the CPR. The ability to manipulate CPR bacteria outside of Saccharibacteria, particularly those with phylogenetically distinct hosts and inhabiting diverse niches, should aid in elucidating the core requirements of the epibiotic lifestyle. Regardless of the precise methods utilized, genetic manipulation of CPR bacteria will open the door to phenotypic studies of the rich reserves of microbial dark matter these organisms contain, potentially revealing unprecedented biological mechanisms.

## Acknowledgements

The authors wish to thank Simon Dove and Joshua Woodward for insightful comments on the manuscript, and members of the Mougous laboratory for helpful suggestions. This work was supported by grants from the NIH (R01AI128215 and R01AI141953 to N.S.B. and R01DE023810 to J.S.M.), the National Science Foundation (MCB-2105570 to N.S.B. and S.T., and IIBR-2042948 and IOS-2050550 to N.S.B.), the Department of Defense, Defense Threat Reduction Agency (HDTRA1-21-1-0007 [DTRA J5B] to D.B.), the Bill and Melinda Gates Foundation (OPP1156262 to D.B.), and the Welch Foundation (I-2095-20220331 to Q.C.). Q.C. is a Southwestern Medical Foundation endowed scholar. J.D.M. and D.B. are HHMI Investigators, and J.D.M. is supported by the Lynn M. and Michael D. Garvey Endowed Chair at the University of Washington.

## Author contributions

L.A.G, A.L., S.B.P and J.D.M conceived the study. Y.W., L.A.G., A.L., I.R.H, S.T., K.J.C., Q.C., N.S.B., S.B.P. and J.D.M. designed the study. Y.W., L.A.G., P.A.A., A.L., and K.J.C. performed experiments. Y.W., L.A.G., I.R.H., S.T., K.J.C., M.L.A-O., Y.L., M.C.R., J.S.M. and Q.C. processed, analyzed and visualized the data. Y.W., L.A.G., I.R.H., S.B.P. and J.D.M wrote the manuscript.D.B., N.S.B., S.B.P. and J.D.M. provided supervision. Q.C., D.B., N.S.B. and J.D.M funded the study. All authors contributed to manuscript editing and support the conclusions.

## Methods

### Strains, media and growth conditions

Saccharibacteria strains employed in this work include *Southlakia epibionticum* ML1 and *Nanosynbacter lyticus* ML1, both isolated in this study. Host bacterial strains used include *Actinomyces israelii* F0345, *A. odontolyticus* F0309*, A. urogenitalis* S6-C4*, Actinomyces sp.* F0386 and *Propionibacterium propionicum* F0230 (*Pp*)^51–53^. Host bacteria mono-cultures, *Se-Ai* co-cultures and *Nl-Pp* co-cultures were routinely grown statically under an atmosphere of ambient air supplemented with 5% CO2 at 37°C or anaerobically with shaking at 37°C using the GasPak EZ Anaerobe Container System with Indicator (BD 260626 and 260001) in TSY media (30 g/L tryptic soy and 5 g/L yeast extract) or TSYR media (TSY media supplemented with 10 mM arginine), or anaerobically on TSBY agar plates (30 g/L tryptic soy, 5 g/L yeast extract, 15 g/L agar, supplemented with 5% (v/v) horse blood). For selection of hygromycin-resistant Saccharibacteria, hygromycin was used at 150 μg/mL. Saccharibacteria-host co-cultures were stored at -80°C in TSY or TSYR supplemented with 10% (v/v) Dimethylsulfoxide (DMSO).

### Isolation of Saccharibacteria strains *Se* ML1 and *Nl* ML1

Isolation of Saccharibacteria strains in co-culture with host bacteria was carried out as essentially as previously described^44, 54^. Anonymous volunteers aged over 18 years provided oral samples for Saccharibacteria isolation. Supragingival plaque samples were collected with toothpicks and dispersed in 1 mL of Maximum Recovery Diluent (MRD; Peptone 1.0 g/L, Sodium Chloride 8.5 g/L, pH 7.0) buffer. 5 mL of saliva was collected by voluntary expectoration into sterile 50 mL conical tubes. All saliva samples and plaque samples were then pooled into 20 mL MRD. Pooled samples were then vigorously resuspended by vortexing and filtered with a 0.22 mm filter, and the flowthrough was collected. Residual bacteria in the sample tube and filter were collected by washing once with 10 mL MRD, filtered again, and combined with the previous flowthrough. Saccharibacteria present in filtrates were pelleted by centrifuging at 60,000 rcf for 1 hr at 4℃. Supernatant was removed, and pellets were resuspended in 1 mL MRD. The presence of Saccharibacteria in these samples were confirmed by PCR, using phylum specific primers^54^ (Table S3). The collected Saccharibacteria were then added to mono-cultures of a panel of five potential host species (*A. odontolyticus, A. urogenitalis, Actinomyces sp.* F0386, *Ai,* and *Pp*) and the cultures were passaged every 24 or 48 hr. The presence of Saccharibacteria cells in the final cultures was confirmed by qPCR with universal Saccharibacteria primers (Table S3) and microscopic imaging. The cultures were then streaked on TSY or TSBY agar, isolated colonies of host bacteria were tested for Saccharibacteria by PCR, and positive colonies were re- cultured in liquid medium and stored as clonal co-cultures.

To create purified suspensions of Saccharibacteria from co-cultures, the co-culture was centrifuged at 3,000 rcf for 5 min at room temperature, to pellet host cells. Supernatant containing suspended Saccharibacteria was then collected and filtered twice using a pre- sterilized 5 μm mixed cellulose esters (MCE) filter. Filtrate was centrifuged at 15,000 rcf for 30 min at room temperature to pellet Saccharibacteria. The resulting pellet was resuspended in a small volume of TSYR supplemented with 10% (v/v) DMSO and stored at -80°C.

### Saccharibacteria host compatibility testing

Purified populations of *Se* and *Nl* for host compatibility testing were generated by growing 100 ml co-cultures of each with their respective host strains according to the standard protocol described above. Co-cultures were passed through a 0.45 μm SFCA filter to remove host cells then spun at 80,000 rcf for 20 min to pellet Saccharibacteria. Supernatant was removed and cell pellets were resuspended in 1 mL fresh medium. Purified Saccharibacteria cells were then add to OD600 = 0.2 cultures of *Ai* and *Pp* (compatible host for *Nl*) at an MOI of 0.2. Cultures were then incubated statically at 37°C under ambient air enriched with 5% CO2 for up to 72 hr, and populations of *Se* and *Nl* were monitored over time using qPCR.

### Whole genome sequencing

Genomic DNA of *Se* ML1, *Nl* ML1and *Ai* F0345 was isolated using the Wizard HMW DNA Extraction kit (Promega). Sequencing was performed on Illumina iSeq and MiSeq and Oxford Nanopore (ONT) MinION instruments after standard sequencing library preparation protocols (Illumina and Oxford Nanopore). De-novo assemblies were generated using the Trycyler pipeline^55^. Specifically, ONT long reads were filtered using Filtlong v0.2.1 (https://github.com/rrwick/Filtlong) with Illumina reference reads, using --keep_percent 95 to retain approximately 95% of the reads. Long reads were then subsampled into 12 bins and assembled into 12 assemblies using the Flye v2.9 (https://github.com/fenderglass/Flye/releases/tag/2.9. Accessed 4 October 2021), Raven v1.8.1^56^ Miniasm v0.3r179^57^ and Minipolish v0.1.3 (https://github.com/rrwick/Minipolish/blob/main/miniasm_and_minipolish.sh) assemblers.

Assembly contigs were manually curated and then reconciled using Trycycler v0.5.2^55^. A consensus assembly for each bacterium was generated and then polished with Illumina short reads using Polypolish v0.5.0^55^ and POLCA from MaSuRCA v4.0.9^58^. Initial annotations were generated using PROKKA v1.14.5^59^.

To sequence clonal transformed *Se* (see Results and below), genomic DNA was isolated from frozen pellets of purified *Se* by re-suspending in Buffer PB (Qiagen) to a total volume of 500 μL, adding 250 μL of 0.1mm zirconia/silica beads (BioSpec Catalog # 11079101z), 250 μL of 20% SDS, and 550 μL of phenol:chloroform:IAA (25:24:1) (Invitrogen Catalog #15593-031), and bead-beating in a Mini-BeadBeater-16 (Biospec Model 607) with settings 3450 RPM, 115V, 10A, and ½ HP, for four 30-second intervals, each followed by cooling on ice for 1 minute.

Purification of the DNA was performed by applying the aqueous phase directly to a DNeasy Blood & Tissue Prep Kit (Qiagen) purification column and following the recommended protocol for washing and elution. Sequencing was performed on an Illumina iSeq using standard library preparation protocols (Illumina). Reads were mapped to the assembled *Se* ML1 genome using minimap2 and variants were called using LoFreq v2.

### Phylogenetic analysis

The *Se* and *Nl* ML1 genomes were phylogenetically placed using whole genome information. A genome tree was generated from these newly isolated strain genomes with a manually curated set of high-quality CPR genomes (complete and partial) and metagenome assembled genomes (MAGS) to remove any contaminants deposited with the original assemblies as described previously^11^. This species tree was constructed using a set of 49 core, universal genes defined by COG (Clusters of Orthologous Groups) gene families with KBase^60^. It combines the curated genome(s) and a set of closely related genomes selected from the public KBase genomes imported from RefSeq. Genome(s) are inserted into curated multiple sequence alignment (MSA) for each COG family. The curated alignments are trimmed using GBLOCKS to remove poorly aligned sections of the MSA. Then, the MSAs are concatenated, and a phylogenetic tree is reconstructed using FastTree2 with the -fastest setting^61^.

### Measuring hygromycin sensitivities of *Ai* and *Se*

Hygromycin sensitivity of *Ai* and *Se* was measured in liquid cocultures. Duplicate co-cultures were initiated by mixing purified *Se* with *Ai* at an OD600 of 0.2 in TSY and at an approximate cellular ratio of 1:2 (*Se*:*Ai*). The co-cultures were divided into multiple aliquots in 96-well culture plates and hygromycin was added to the final concentrations shown in Figure S1. The plates were covered with Breath-Easy sealing membrane (Sigma Z380059) and incubated without lids at 37°C with 5% CO2. At 1 day and 3 days, the cells within individual wells were pelleted (>15 min at >15,000 rcf) and stored at -20°C for later genomic DNA isolation and qPCR-based quantification of *Se* and *Ai* (see below).

### Design and generation of cassettes for heterologous gene expression in *Se* and *Nl*

Cassettes for heterologous gene expression in *Se* and *Nl* were designed by appending promoter and terminator sequences from the TM7x genome^8^ to the 5’ and 3’ ends, respectively, of ORFs codon optimized for *Se*. The promoter and terminator elements were sourced from TM7x rather than *Se* to reduce the likelihood of off-target integration at corresponding *Se* loci. The TM7x promoters were chosen from genes expected to be highly and constitutively expressed, *tuf* and *rpsJ*. *Se* codon usage was calculated using the Dynamic Codon Biaser (DCB)^62^. Heterologous ORFs were optimized to match relative codon usage frequencies found in *Se,* but omitting codons with less than 10% usage. Heterologous genes utilized included *hph* (hygromycin B phosphotransferase) from *Streptomyces hygroscopicus*, superfolder GFP (www.fpbase.org/protein/superfolder-gfp/), mCherry2 (www.fpbase.org/protein/mcherry2/) and NanoLuc Luciferase^63, 64^. Table S4 reports the composition and complete sequences of the designed cassettes. Cassettes were obtained as gBlocks from Integrated DNA Technologies, Inc (IDT).

Linear fragments used for transformations were generated by adding *Se* or *Nl* genomic sequences corresponding to the targeted insertion or allelic replacement sites to the left and right sides of a heterologous gene expression cassette or of two cassettes joined together. Overlap extension PCR was used to join gBlock cassettes and genomic fragments, the latter of which were individually amplified from *Se* or *Nl* genomic DNA. In some cases, complete fragments including genomic sequences were obtained as gBlocks from IDT. Fragments used for flank- length tests were generated by amplification using larger fragments as templates followed by gel-purification. Table S4 reports the composition and complete sequences of the fragments utilized and primers used are listed in Table S3.

### Genetic transformation of *Se* and *Nl* with targeted insertion and allelic replacement constructs

To prepare *Se*-*Ai* or *Nl-Pp* co-cultures for transformation, 1-mL aliquots of a previously frozen co-culture (see methods on co-culturing) were thawed on ice and added to 9 mL of TSY supplemented with freshly cultured *Ai* to a final OD600 of 0.2, incubated statically for 2 d at 37°C in ambient air enriched with 5% CO2, then enlarged by the addition of 10 volumes of TSY and incubated anaerobically using the GasPak EZ Anaerobe Container System with Indicator (BD 260626 and 260001) at 37°C for 2 days with shaking at 160 rpm.

For transformation of *Se* or *Nl* with linear targeted insertion constructs, 0.22 - 0.3 mL aliquots of prepared co-culture were incubated statically with transforming DNA for 6 h at 37°C in ambient air enriched with 5% CO2 in culture tubes. The mixtures were subsequently enlarged by the addition of TSYR to approximately 5 mL and supplemented with recently passaged *Ai* (for *Se*) or *Pp* (for *Nl*) to a final OD600 of approximately 0.06. This time point was designated “TF” (time zero after transformation). After removal of a sample for later qPCR analysis, the cultures were divided and one half was supplemented with hygromycin to 150 μg/ml. The cultures were then incubated as above for an additional 2-3 days. This time point was designated “P1” (end of initial passage). Cultures were subsequently serially passaged as many as four times by five-fold dilution into fresh TSYR supplemented with *Ai* (for *Se*) or *Pp* (for *Nl*) to a final OD600 of 0.06 and, for the hygromycin containing cultures, with additional hygromycin to 150 μg/ml. These passages (designated “P2”, “P3”, etc.) were each incubated for 2-3 days at 37°C in ambient air enriched with 5% CO2 and without agitation. Cultures were sampled at each passage by removing and pelleting (>15 min at >15,000 rcf) of 0.1 to 1 ml, and the pellets were stored at -20°C for later qPCR analysis.

### qPCR assays

We employed qPCR for quantification of Saccharibacteria and host populations in cocultures, as well as to monitor *Se* transformation. For these assays, genomic DNA was isolated from frozen co-culture pellets using Instagene matrix (Bio-Rad) and quantitative PCR (qPCR) was performed using a CFX Connect Real-Time PCR Detection System (Bio-Rad). To quantify *Se* and *Ai*, amplification employed primers targeting *Se uvrB* or the 16S rRNA gene of *Ai* and was performed in 20 μL reactions with 1X SsoAdvanced Universal SYBR Green Supermix (Bio-Rad), 300 nM each primer and 4 μL template DNA (primer sequences provided in Table S3). Thermocycling conditions were 95°C for 5 min followed by 35-40 cycles of 95°C for 20 s, 60°C for 30 s and a read of fluorescence. To quantify transformed *Se* (insertions at NS1), amplification employed primer pairs targeting the insertion sequence and sequence adjacent to NS1 but outside of the linear transformation construct arms and was performed in 20 μL reactions with 1X Phusion HF Buffer, 0.2 mM dNTPs, 250 nM each primer, 0.5X SYBR Green I (ThermoFisher), 4 μL template DNA and 0.4 units of Phusion High-Fidelity DNA Polymerase (NEB) (primer sequences provided in Table S3). Thermocycling conditions were 98°C for 90 s followed by 35- 40 cycles of 98°C for 15 s, 70°C for 20 s, 72°C for 90 s and a read of fluorescence. Melt curves were performed following each amplification to verify product homogeneity. Absolute target sequence abundance was determined by comparison to duplicate standard curve reactions performed in parallel with each assay. The standard curve templates were generated by serial dilution of previously amplified and gel-purified products quantified by Qubit (ThermoFisher).

### Isolation of isogenic mutant co-cultures

To obtain co-cultures with isogenic mutant (transformed) *Se*, transformation mixtures grown for three passages under hygromycin selection were serially diluted, plated on TSBY agar without antibiotic and incubated for 7 days anaerobically. Isolated colonies (representing *Ai* potentially colonized with *Se*) were picked into 0.2 mL TSYR supplemented with *Ai* at a final OD600 of 0.2 and incubated statically at 37C in ambient air enriched with 5% CO2 for 2-4 days, then screened for the presence of *Se* and for the mutant (genomic insertion) and wild-type (no insertion) alleles by PCR (e.g., Figure 2A, primer sequences in Table S3). Cultures containing pure mutant *Se* were frozen and/or further propagated for purification of the *Se* for WGS (see above).

### Nanoluciferase assay

Nanoluciferase assays were performed using isogenic wild-type *Se-Ai* co-cultures and *Se* NS1::*nluc-hph2-Ai* co-cultures grown statically in ambient air supplemented with 5% CO2 at 37°C in TSY media for three passages as described above. To separate *Se* from host cells, 20 mL of each co-culture was passed through a 0.45 μM SCFM filter and the resulting filtrate was spun at 80,000 rcf for 30 min to pellet *Se*, supernatant was removed, and the pelleted *Se* was resuspended in 750 μL TSY. 100 μL of these purified *Se* cells or 100 μL of the corresponding *Se-Ai* cocultures were mixed with 100 μL Nano-Glo Luciferase assay reagent (50:1 mixture of substrate:buffer, Promega N1110) in a 96-well plate. Luminescence signal indicative of nanoluciferase activity was detected using a Cytation 2 plate reader. Luminescence signal was later normalized by *Se* abundance in co-cultures and filtrates as measured by qPCR using *Se*- specific primers as described above.

### Microscopy

Imaging was performed on a Nikon Eclipse Ti-E wide-field epi-fluorescence microscope, equipped with a sCMOS camera (Hamamatsu) and X-cite LED for fluorescence imaging. We imaged through a Nikon Plan Apo λ 60X 1.4 NA oil-immersion Ph3 objective. The microscope was controlled by NIS-Elements 3.30.02. Isogenic *Se* NS1::*mcherry-hph2-Ai* co-cultures and *Se* NS1::*sfgfp-hph2-Ai* co-cultures were grown statically in ambient air supplemented with 5% CO2 at 37°C in TSYR media as described above. Cell samples were spotted on a 3% (w/v) agarose pad made with TSYR media supplemented with 0.4% glucose placed on a microscope slide. The microscope chamber was heated to 37°C for time-lapse experiments. Time-lapse images were aligned and segmented with Omnipose using the phase contrast channel and the published bact_phase_omni model^65^. Masks were manually linked and corrected for segmentation errors in regions of *Ai* cell overlap using Napari. Regions corresponding to *Se* were also removed to accurately track host *Ai* growth alone. For visualization purposes, fluorescence intensity was gamma-corrected to normalize background levels on a frame-by-frame basis while not distorting the *Se* signal. No bleaching correction was implemented. Figure 3A-F and Videos [S1-11](https://youtube.com/playlist?list=PLSQI8BpslH-PhSKoxXLm7Ur8zJneoS5We) were generated using Python and annotated in Adobe Premiere.

### Mutagenesis of Sac1a genomic DNA and transposon mutant library generation

The transposon used for *in vitro* transposition (here named T36) was generated by amplification of the *hph1* insertion cassette using the 5’-phosphorylated primers T36-ampF and T36-ampR, which add required 19-bp Tn*5* mosaic end sequences to each end of the amplicon, followed by purification using the Qiagen PCR Purification Kit with elution in TE (primer sequences provided in Table S3). *In vitro* transposition was performed in multiple 45-μl reactions containing 1.8 μg *Se* genomic DNA, 290 ng purified T36, 1X EZ-Tn5 Reaction Buffer and 1.2 U EZ-Tn*5* Transposase (Biosearch Technologies) with incubation for 2 hr at 37 °C. The reactions were stopped, DNA was ethanol precipitated, and gaps were repaired as described^66^.

Following gap repair, the DNA was purified using Qiagen PCR Purification Kit with elution in water after the columns were washed twice with Buffer PE.

Transformation of *Se* to generate a transposon mutant library was performed similarly to the procedure described above for transformation with targeted insertion fragments, but at a larger scale and with modifications. Specifically, the initial 6 hr incubation contained 4.0 μg of *in vitro* transposon-mutagenized genomic DNA and 15 mL of *Ai*-*Se* co-culture. This co- culture was enlarged by adding 135 mL of TSYR supplemented with *Ai*, then incubated for two additional hours before hygromycin addition. Concomitantly with hygromycin addition, MgCl2 (to 1 mM) and benzonase (Sigma) (to 25 U/mL) were added to degrade extracellular DNA, and the cultures were similarly supplemented with benzonase approximately every 24 h during passaging. After the initial 2-day growth passage (ending at time point T0), the culture was serially passaged four additional times by dilution of 30 mL culture into 120 mL in TSYR supplemented with *Ai* at OD600 of 0.067. These passages were each incubated statically for 2 days at 37°C in ambient air enriched with 5% CO2. Samples for Tn-seq analysis were taken at the end of the initial passage (60 mL, T0) and at the end passages 1, 2 and 4 (80 mL, T1-T3).

These samples were treated with additional benzonase (50 U/mL) for 30 min, then EDTA was added to 15 mM and pellets were collected by centrifugation for 30 min at 15,000 rcf and stored at -80°C.

### Tn-seq library preparation and sequencing

Genomic DNA was isolated from frozen co-culture pellets using the bead-beating method described above (section on whole genome sequencing). Tn-seq libraries were prepared by the C-Tailing method as described^67^. The transposon-specific primers utilized are listed in Table S3. Sequencing was performed in multiplex as 50-bp single-end reads on an Illumina MiniSeq with 25-40% PhiX spike-in.

Custom scripts (https://github.com/lg9/Tn-seq66,67) were used to process the Illumina reads. In brief, reads were first filtered for those displaying transposon end sequence as their initial bases (the sequencing primer was designed to anneal six bases from the end of the transposon). These reads were then mapped to the *Se* genome after removing the transposon end sequences. Reads per unique mapping position and orientation were tallied and read counts per gene were calculated by summing reads from all unique sites within a given gene.

### *Se* gene fitness and essentiality analysis

We first checked the saturation level (i.e., number of insertions detected) of the Tn-seq library sampled along four time points (T0-T3). Because the T0 sample had the highest number of unique insertions (9,996), we initially focused on this time point to identify essential genes in the *Se* genome. Gene essentiality was determined based on the number of insertions detected in each gene (using the GUMBEL method, part of the TRANSIT tool), or alternatively by taking into account the read counts per insertion(s) in each gene (using the Hidden Markov Model method, HMM, also part of TRANSIT)^26^. Default parameters were used for both GUMBEL and HMM analyses. To complement these analyses and to identify genes whose disruption was deleterious for *Se* in later time points, we used a comparative approach with respect to T0. The comparative approach was necessary due to the significant reduction in the number of unique insertions detected at T1-T3 (the average number of insertions for these time points was 1,765), which suggested a strong selection pressure over time. Thus, the distribution of insertions at T1, T2 or T3 were independently compared with T0. We detected genes associated with differentially abundant insertions at T1-T3 vs. T0 using TRANSIT and ALDEx2^26, 27^. For the TRANSIT-based analysis, we used the Resampling method to identify differentially abundant genes between the two time points being compared. Genes with TRANSIT-estimated adjusted p-value < 0.05 were considered as differentially abundant (this gene set would include under- and over-represented genes at the late time points with respect to T0). ALDEx2 takes into account the compositional nature of sequencing data generated during selection assays (i.e., competitive growth mutant assays)^27^. Given the lack of replicates (expected in the standard ALDEx2 pipeline) in our data, we adapted the ALDEx2 workflow by using the aldex.clr function available in the ALDEx2 R package to generate Dirichlet distributions of centered-log ratio (CLR) transformed abundances for any pair of samples being compared (e.g., T1 vs. T0)^27^. We used 1,000 Monte Carlo instances to generate the Dirichlet distributions. Then, we computed the median CLR value per gene for each sample. Finally, we computed a delta score similar to ALDEx2 original “dif.btw” by computing gene wise median difference between the two samples being compared (e.g., T1-T0). Genes with delta scores < -1.5 were considered under-represented in the late time points (with respect to T0), suggesting that insertions that disrupt those genes were deleterious for *Se*. To define a meaningful threshold for our delta score and confirm the performance of our ALDEx2-based analysis, we leveraged barcode sequencing data we previously generated for competition growth assays of an antibiotic-treated *Escherichia coli* pooled single-gene deletion library^68^. This benchmarking dataset included four time points (T0- T3), each one with four replicates. For the T1 vs. T0 and T2 vs. T0 comparisons we applied our strategy for each individual replicate and then evaluated the overlap with the original results obtained when considering all four replicates. We found that indeed the set of genes with absolute delta score > 1.5 for single replicate analyses significantly overlap with the set of genes in the analysis that included all replicates. Three out of the four single replicate analyses had hypergeometric test p-values < 0.05 for T1 vs. T0 (two with p-values < 0.005). Moreover, all single replicate analyses for T2 vs T0 had hypergeometric test p-values < 1E-202.

### Transformations to validate gene essentiality predictions

To validate Tn-seq essentiality predictions, we designed constructs to replace target genes with *hph.* Each construct contained 335-bp sequences targeting the desired site flanking the *hph* expression cassette described above. 100 ng synthesize linear DNA or water were added to 300 mL wild-type *Se-Ai* co-cultures (4 biological replicates per condition) prepared according to our standard protocol, and transformation and subsequent passaging were performed using our general transformation protocol described above. At the end of passage 3, 1 mL of each transformed co-culture was treated with 1 mM MgCl2 and 25 U/mL benzonase for 30 min at 37℃ to remove extracellular DNA. 15 mM EDTA was then added to stop the benzonase reaction, samples were centrifuged at 21,000 rcf for 1 hr and the resulting pellets were stored at - 80℃. *Se* abundance was then measured by qPCR as described above.

### Generation and transformation efficiency of *Se* /-*comEC::hph*

A clonal population of *Se* /-*comEC::hph* was generated by replacing *comEC* with our hygromycin resistance cassette following the transformation and clonal mutant isolation protocols described above. To assess *Se* /-*comEC::hph* transformability, 50 ng of synthesized linear gene product for inserting unmarked *sfGFP* at NS1 (with 545 bp flanking regions) was used to transform 150 μL isogenic wild-type or /-*comEC Se-Ai* co-cultures (4 biological replicates per condition). After 6.5 h of incubation between DNA and co-cultures at 37℃, the entire transformation culture was treated with 1 mM MgCl2 and 25 U/mL benzonase for 30 min at 37℃ to degrade extracellular DNA. 15 mM EDTA was added to stop the benzonase reaction and the samples were centrifuged at 21,000 rcf for 1 hr and the resulting pellets were stored at -80℃. We then used qPCR to measure the abundance of transformed and total *Se* using NS1- sfGFP or *uvrB-* specific primers as described above.

### CPR-enriched protein families analysis

To assess the protein family distribution of the *Se* and *Nl* genomes, we collected amino acid sequences for 22,977 protein clusters (protein families) from Meheust *et al*.^5^. These protein families were built through their occurrences across at least 5 distinct non-redundant or draft CPR bacteria, non-CPR bacteria and Archaeal genomes. The presence or absence of any given family in *Se* or *Nl* were determined through all-vs-all protein sequence search of every protein in the genome against the database of protein family sequences. Protein sequence search was performed by using the MMseqs2 (version: 14-7e284) algorithm with the following parameters: module: easy-search; sensitivity (-s): 7.5; alignment coverage (-c): 0.5; greedy-best-hits: 1. The resulting presence or absence matrix was combined with the core family matrix of 921 protein families across 2890 genomes (collected from Meheust *et al*.^5^) for clustering analysis. A Jaccard distance based on the presence or absence of core families across *Se*, *Nl* and 2890 other genomes was calculated in R by using proxyC package (version: 0.3.3). Agglomerative hierarchical clustering was performed by using cluster package (version: 2.1.4) in R with the “wand” method. A hierarchical clustering heatmap was built using complexHeatmap package in R (version: 2.14.0) by plotting presence (black) or absence (white) of protein families (columns) in a given genome (rows). Enrichment or depletion of core protein family clusters in *Se* and *Nl* was performed using Fisher’s exact test in R with input table of presence or absence of protein families in the *Se* and *Nl* genomes. A threshold of Benjamini-Hochberg corrected p-values of < 5e-05 was used to determine the enrichment or depletion.

### Sequence-based gene annotation of the *Se* genome

We used ProtNLM (https://github.com/google-research/google-research/tree/master/protnlm) to predict the function annotations of proteins in the *Se* proteome from sequence^30^. A prediction of score ≥0.5 corresponds to >70% accuracy for proteins without close UniRef50 matches and ∼95% accuracy for proteins with close matches. We thus considered the 337 proteins with score ≥0.5 to have confident annotations (Table S2).

Additionally, we mapped each protein to Pfam domains (Nov, 2021 release v35.0^46^) using hmmscan^31^. We filtered the hit Pfam domains by full sequence E-value ≤ 1e-3 and domain-based CE-value ≤ 1e-5 or CI-value ≤ 1e-5, domain coverage ≥ 50% (coverage of aligned portion of domain to full length domain) and selected the top hit for each aligned region.

### Generation of multiple sequence alignments

To construct MSAs for each protein in the *Se* proteome, we initially used HHblits^35^ to search against UniRef (2022 release^37^) and BFD (2019 release^36^). For those MSA that contained <500 sequences after this approach, we conducted extensive homology searches against metagenome databases similar to Anishchenko *et al*.^69^. We first converted the initial HHblits (E-value ≤ 1e-3) MSAs for each protein into a hidden Markov model (HMM) which we used as a seed sequence profile to search against metagenomic and metatranscriptomic sequences from JGI^70^, MGnify^71^, and UniRef with hmmsearch^31^, respectively. Hits from JGI, MGnify, and UniRef were gathered and aligned to the query protein by phmmer and jackhammer^31^, to retain sequences with E-value ≤ 1e-5 at each stage. We removed columns containing gaps in the query sequence from the MSAs, which were then subjected to redundancy filtering at 95% sequence identity and 50% sequence coverage. We selected the deepest MSA for each protein (from HHblits against UniRef+BFD or phmmer/jackhmmer against metagenome) for downstream analysis.

### Structure-based annotations

Using MSAs described above, we computed AlphaFold (AF) models (model_1 without structural template search, 10 recycles, and version 1.0 weights) for 852 of the 855 proteins (3 were excluded because of MSA depth or protein size)^34^. We then applied FoldSeek^38^ against AFdatabase50 (AF Protein Structure Database clustered at 50% sequence identity and 80% coverage). We report the top hits in Table S2 based on E-value ≤ 0.00001, TM-alignment score ≥ 0.5 (which correlates well to fold-family level similarity^51^), and query coverage and target coverage ≥ 50%. To classify proteins into evolutionary contexts based on structure and sequence similarity, we used Domain Parser for AlphaFold Models (DPAM)^42^, and the full results are available at https://conglab.swmed.edu/ECOD_Se/Se_ECOD.html.

### Nucleotide sequence accession numbers

The complete genome sequences of *Se* ML1, *Nl* ML1and *Ai* F0345 have been deposited in GenBank under BioProject PRJNA957798 with accession numbers SAMN34266291, SAMN34266292 and SAMN34266293, respectively. Tn-seq data generated in this study have been deposited in the Sequence Reads Archive (SRA, BioProject PRJNA957798).

## Supplemental information

**Figure S1.**
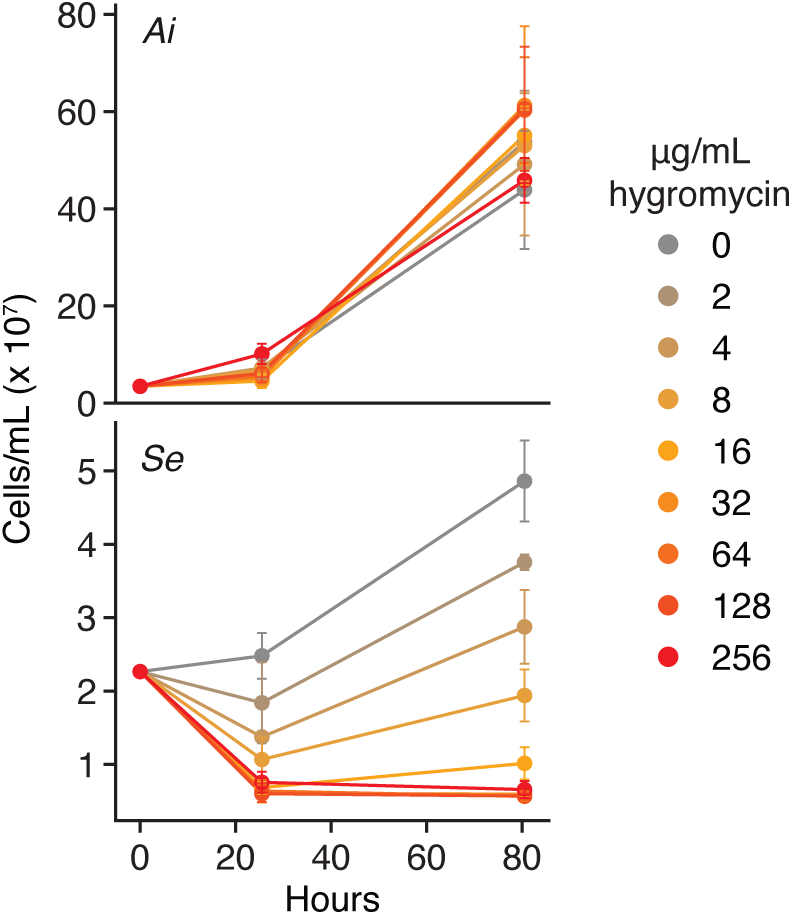
Hygromycin resistance phenotypes of *Se* and *Ai* enable use of *hph* as a selectable marker in *Se.* Growth of *Ai* (top) and *Se* (bottom) in co-cultures containing the indicated concentrations of hygromycin.

**Figure S2.**
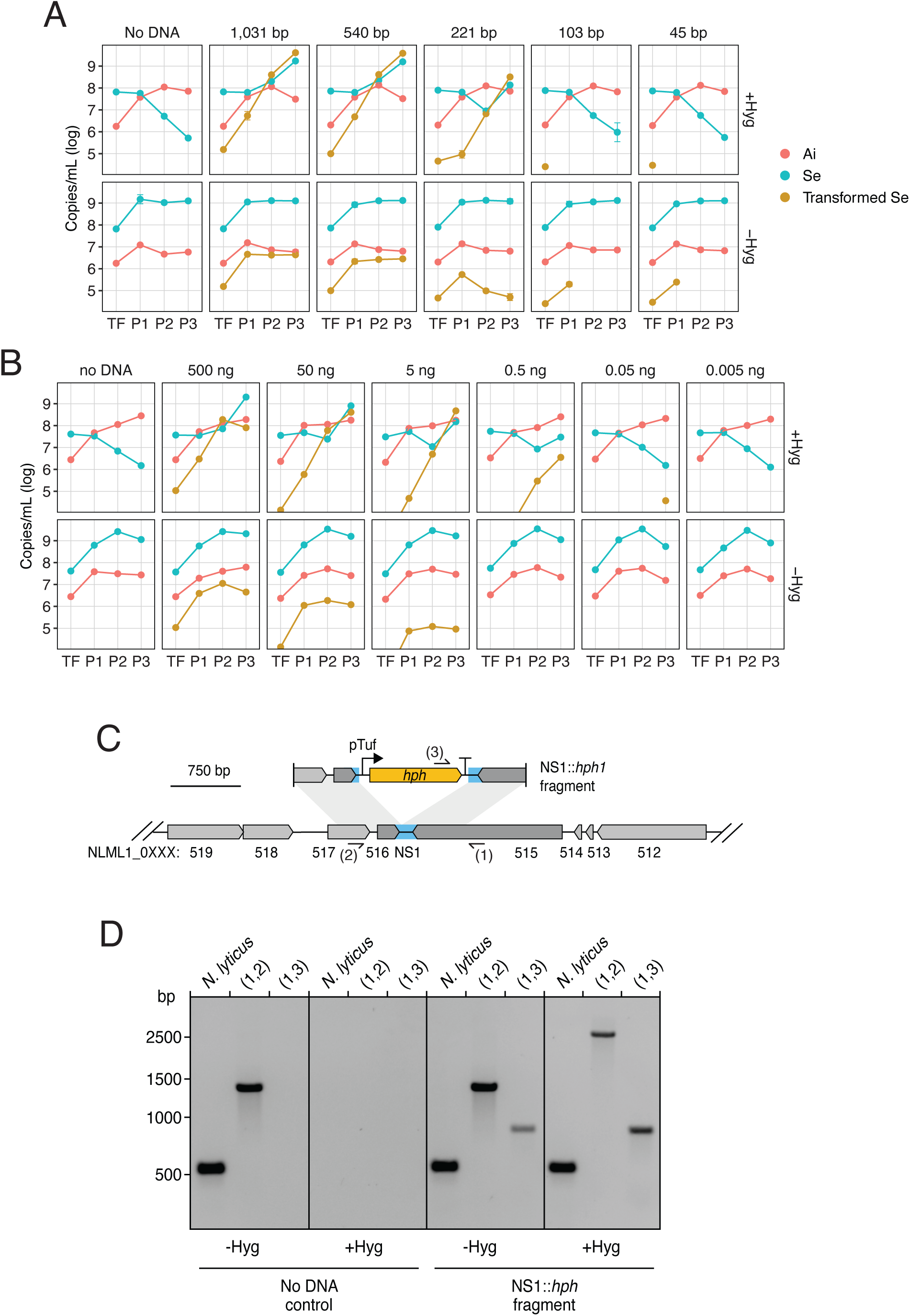
Optimization of the *Se* transformation protocol and transformation of a second Saccharibacteria species. (A, B) Quantification of *Se, Ai* and transformed *Se* populations over the course of our transformation protocol (See Figure 2B) with varying lengths (A) or concentrations (B) of transforming DNA. Transformations were performed in the presence (top panels) and absence (bottom panels) of selection for transformants with hygromycin. (C) Schematic depicting the intergenic neutral site (blue, NS1) targeted for insertion of a hygomycin resistance cassette (yellow) in the *N. lyticus* ML1 genome and the linear DNA fragment employed in transformation experiments with this species. Primer binding sites used for genotyping are indicated (sites 1-3). (D) Genotyping of *N. lyticus–P. propionicum* co-cultures transformed with the linear DNA fragment depicted in (C) (right) or parallel negative control co-cultures with no DNA added (left) at the end of passage 4 (see Methods) with primers targeting NS1 (1,2) or *hph* (3). Positive control *N. lyticus* primers target a genomic locus distant from NS1.

**Figure S3.**
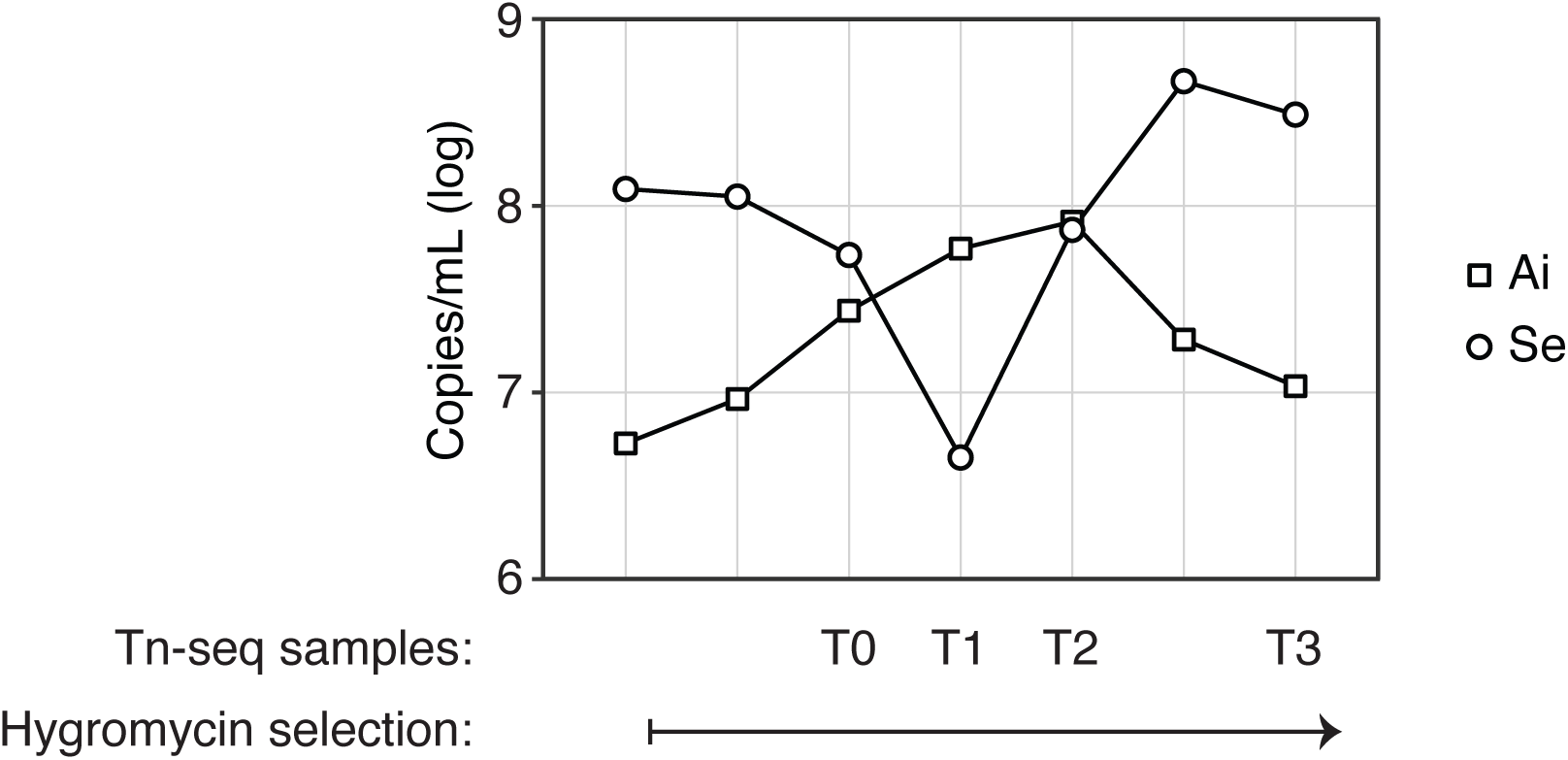
Population dynamics of *Se* and *Ai* during transposon mutagenesis. Transformation co-culture was diluted and passaged into fresh media, with the addition of fresh *Ai,* after collection of the T0, T1 and T2 samples (see Methods).

**Figure S4.**
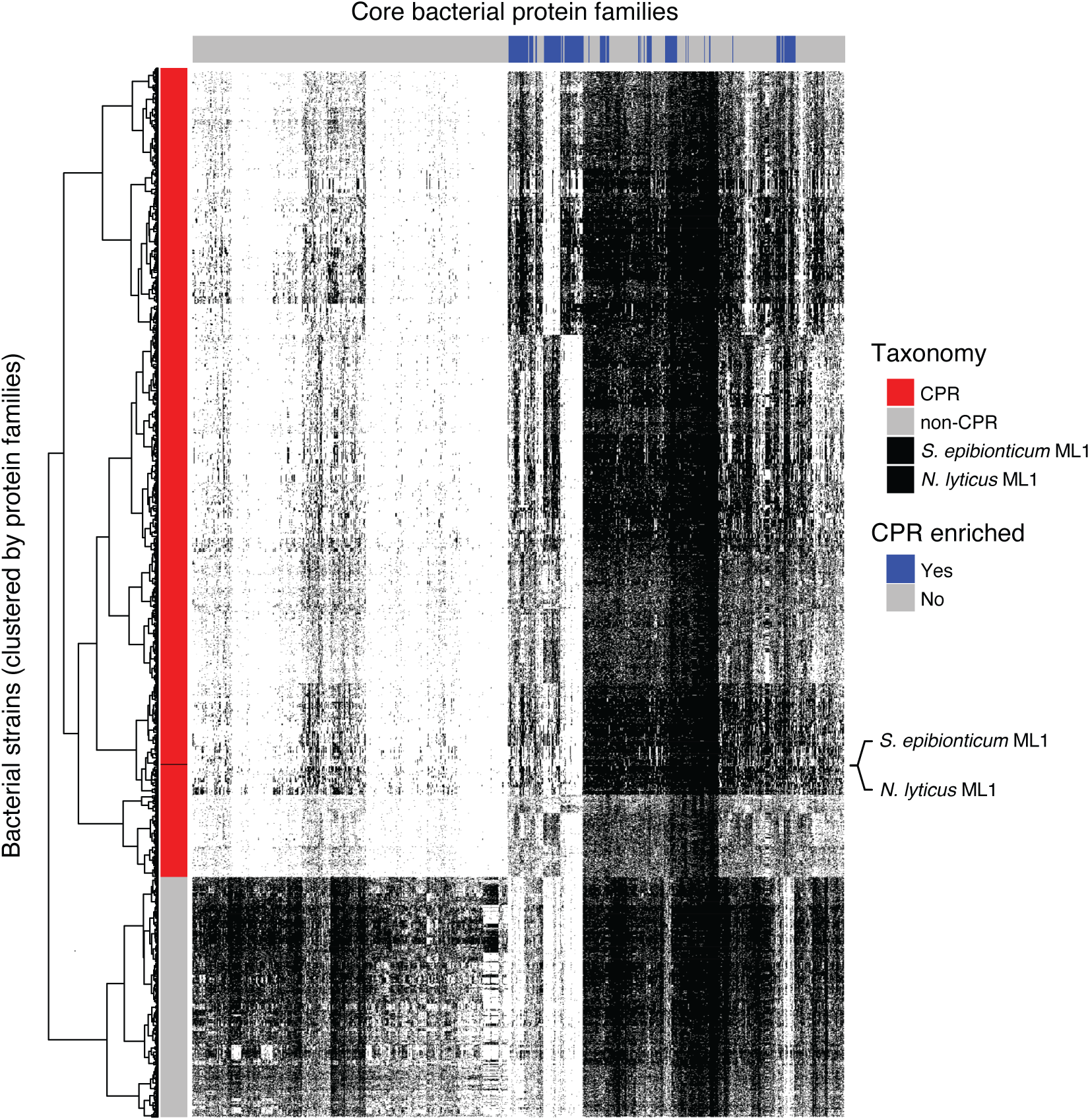
Distribution of 921 core protein families across CPR and non-CPR bacterial genomes, including *Se* and *N. lyticus* ML1. Columns represent core families (derived from Meheust *et al.*^5^) and rows represent individual genomes from the indicated bacterial groups. CPR enriched protein families indicated at top (blue), and dendrogram at left represents clustering of bacterial strains based on protein family content.

**Figure S5.**
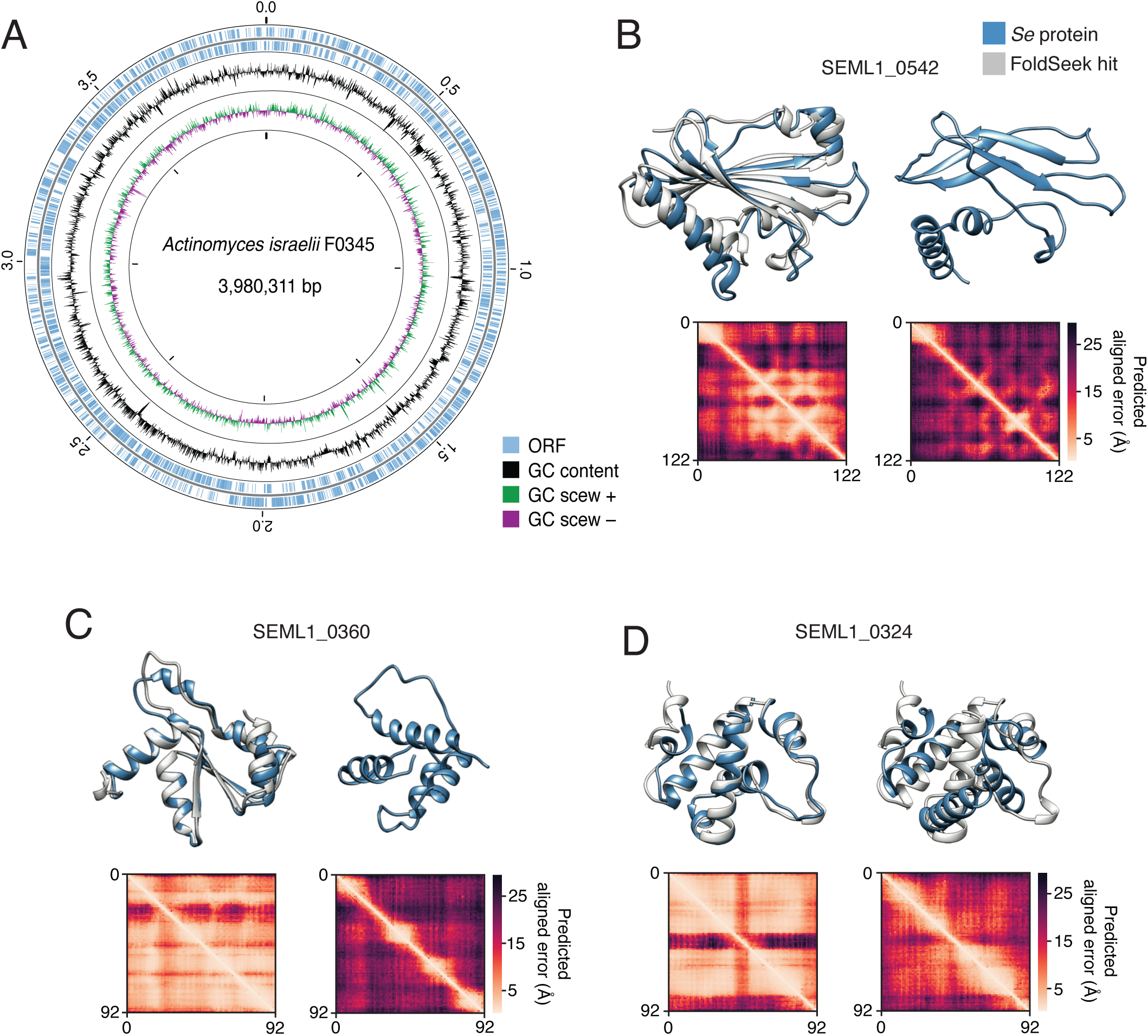
The *Ai* genome and structural models for the *Se* proteome generated using metagenomic sequence enriched MSAs for use in future studies of the *Se-Ai* system. (A) Overview of the genome sequence of *A. israelii* F0345. (B-D) Example *Se* protein structure models and associated predicted alignment matrices obtained using shallow (right) or metagenomic sequence-improved (left) MSAs. *Se* proteins models (blue) are aligned to models for top FS hits (light grey, AF database50 numbers A0A8B1YQG7 (B), A0A660LZS9 (C) and A0A563CGZ5 (D)), when available. Some structures are trimmed to highlight the alignment.

Table S1. Tn-seq data and analysis. Shaded rows indicate genes found to be important for *Se* fitness during in vitro co-culture with *Ai* by both TRANSIT and ALDEx2, at all time points sampled. Blue text indicates genes within loci noted at the perimeter of Figure 4A.

Table S2. Sequence and structure-based annotation of the *Se* genome. Shaded rows indicate genes encoding proteins for which the AlphaFold confidence score (pLDDT) was improved by 10 or more points upon inclusion of extensive metagenomic data in the MSA.

Table S3. Oligonucleotide primers employed in this study.

Table S4. Heterologous gene expression cassettes and linear fragments used in Saccharibacteria transformations.

